# Machine Learning and Deep Learning challenges for building 2′O site prediction

**DOI:** 10.1101/2020.05.10.087189

**Authors:** Milad Mostavi, Yufei Huang

## Abstract

2′-O-methylation (2′O) is one of the abundant post-transcriptional RNA modifications which can be found in all types of RNA. Detection and functional analysis of 2′O methylation have become challenging problems for biologists ever since its discovery. This paper addresses computational challenges for building Machine Learning and Deep Learning models for predicting 2′O sites. In particular, the impact of sequence length containing 2′O site, embedding method and the type of predictive model are each investigated separately. 30 different predictive models are built and each showed the impact of the mentioned parameters. The area under the precision-recall and receiving operating characteristics curves are utilized to test imbalanced case scenarios in the real world. By comparing the performance of these models, it is shown that embedding methods are crucial for Machine Learning models. However, they do not improve the performance of Deep Learning models. Furthermore, the best predictive model was further investigated to extract significant nucleotides surrounding 2′O sites. Interestingly, based on the significant score matrix achieved by all 2′O samples, it is depicted that model pays the highest attention at the location that the dominant 2′O motifs exist. Dataset and all of the codes are available at https://github.com/MMostavi/2_O_Me_sitePred

## I. Introduction

There are more than 150 different chemical RNA modifications that influence the coding information of RNA. 2′O is one of the major modifications that occur in different RNA types in all domains of life. This modification can be formed in all four types of RNA nucleotides of A (adenine), C (cytosine), G (guanine), and U (uracil). As a result, this intrinsic behavior makes the precise detection and functional analysis of this RNA modification a challenging task. Several research studies have focused on detecting the location of 2′O in different species, whereas other papers have investigated the specific functional effect of 2′O, such as splicing [1].

Deep Learning (DL) models have shown unprecedented success in a wide variety of applications ranging from classifying human brain signals to predicting sites of epitranscripome modifications [2, 3]. In particular, Convolutional Neural Networks (CNNs) can be extremely beneficial in biological datasets not only by achieving high accuracy in predictive tasks but also by providing a clear explanation of the inner working process of the model [4, 5]. Furthermore, automatic feature extraction in DL models is eliminating conventional human-made feature extraction methods that require a well-established prior knowledge about the biological problem of interest. This paper is an extension of our previous work in [6] to further investigate modeling and interpretation of 2′O methylation sites.

In this paper, first, we will investigate three important key factors in building models for 2′O methylation site prediction: length of 2′O input sequences, and impact of different DNA/RNA embeddings in DL and ML predictive models. In particular, the impact of each factor is evaluated by the performance of predictive models. One-hot encoding and dna2vec are selected as two of the candidate embedding methods which are used in most ML and DL sequence-based models. For predictive models, Support Vector Machines (SVMs) and CNNs are used as the best ML and DL predictive models for sequence-based problems. Once the model is established, we will dive into the interpretation of the best model to extract the importance of neighboring nucleotides. Specifically, we are interested to investigate what the significant nucleotides are from a model perspective during prediction.

The organization of the paper is as follows: in the next section, the dataset that is used followed by an explanation of two embedding methods and predictive models are described. In the results section, first, we will show the performance of all models. Second, we select one of the models and delve into the interpretation of the model in order to understand the working process of the model.

## II. Materials and Method

### A. Dataset

The locations of all 2′O methylation sites for H. sapiens were collected from nm-seq technology in [7] and aligned with hg38 human genome with BedTools to extract surrounding sequences. There are 2802 locations in two cell lines, HEK and HeLa 699 and 2103 respectively that represent 2′O sites in all four RNA nucleotides. Furthermore, 7004 sequences were randomly selected as negative samples through the whole human transcriptome if they did not contain any 2′O sites.

600 of 2802 positive samples (i.e. around 20 percent) and 3000 of 7004 negative samples were held out for testing where remaining samples were allocated for training models. A ratio of 1:2 was used in training.

#### Length of input sequences

According to biologists, the flanking region of site of interest has to be long enough to include most expected motifs and at the same time short enough to exclude non-motif occurrence site positions [8]. From a computational standpoint, the former invigorates the power of predictive model and latter generates noise that need to be handled. In order to delineate the impact of each side of 2′O modification sites and the length of surrounding nucleotides, we hypothesis two different scenarios for choosing the best input sequence.

First, like many sequence related papers [3], we assume upstream and downstream sides of 2′O sites have same contribution in its occurrence. In this case, the origin of interest (i.e. 2′O site) is anchored in the middle of surrounding nucleotides in a symmetry fashion and the length of sequence (L) varies from 45 nucleotides to 75 nucleotides, and 105 nucleotides to observe the impact of sequence length. In the embedding section, we describe why these specific lengths are selected.

Second, 2′O motif search in [7] suggests that 2′O sites have a dominant motif in their downstream known as N_m_AGAUCGAGGA where N_m_ can be any of four RNA nucleotides. Computationally speaking, this motif can be interpreted as a fixed pattern that leads to reaching better performance results when downstream nucleotides of 2′O site are used for training predictive model compared to its upstream. In this case, 2′O sites with either their downstream or upstream side were selected with a window size of 44 nucleotides.

### B. Embedding

The human genome is described by four distinct letters where their assortment primes human genetic code. A substantial amount of research studies are devoted to finding the best practices for embedding sequences into useful representation, which later can be used in predictive models [9, 10]. Hereby, we attempt to exploit one-hot-encoding as the widely used representation method and dna2vec as DL based embedding method.

#### One-hot encoding

One hot encoding takes categorical variables such as A, C, G, U in the case of RNA sequences and implicitly converts them to numerical variables to increase the ordinal properties of categorical variables for a predictive model. These numerical values are mostly represented as columns of Boolean numbers, where the value of one is assigned to the presence of one category in the corresponding column. This concept is shown in **Figure 1**.

**Figure 1:**
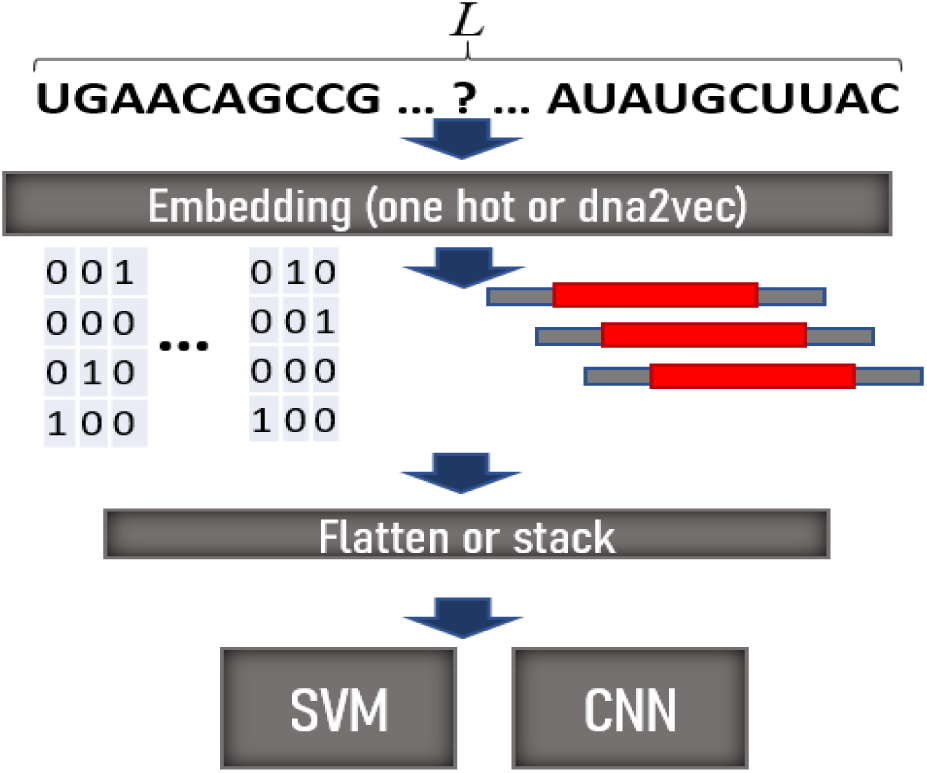
Scheme for investigating input sequence length, embedding methods, and predictive models.

Although one hot encoding has a straightforward and simple formula for embedding RNA sequences, it has major drawback that limits its usage. This embedding suffers from the curse of dimensionality. meaning that by increasing the length of RNA sequence, the binary feature representation grows exponentially. This fact leads to low-performance results in ML models that require low dimension inputs.

#### dna2vec embedding

This method converts sequences with length kmer where *kmer* ∈ {3,8} to fixed vectors with length 100 [11]. The biggest advantage of dna2vec is the ability to capture arithmetic relations between input sequences.

2′O and non-2′O sequences are embedded with kmer equal to three and five as described in [6]. Specific input lengths were a factor of multiplied these two kmers. Although the output vector length of dna2vec is fixed, variable kmer gives us this advantage that we can observe the impact of input sequence alterations.

### C. Predictive models

CNN and SVM are two of the predictive models that we used to investigate the effect of different embeddings and input sequence length on their performance. The main reason that we benchmark SVM among different ML models is because several papers have reported that SVM has had superior performance over other ML models in sequence-based problems [12].

As it is shown in **Figure 1**, a sequence with length L will be embedded either with one-hot or dna2vec. Depending on the predictive model in the next step, the output of embedding will be reshaped either into an image-like input (to be used in CNN) or a flattened vector (to be used in SVM). One hot encoding generates matrices with 4*L dimension for every sequence. On the other hand, dna2vec generates 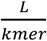 ∗ 100 inputs for every sequence. For instance, if L = 45 and kmer = 3, dna2vec will generate 15 vectors with length 100. If SVM is used, these vectors will be stacked horizontally otherwise they will be stacked vertically to create matrices with 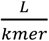 ∗ 100 dimension. In [6, 13] further details on the implementation of CNN models are described.

In order to evaluate the performance of predictive models, the area under Precision-Recall (auPR) and area under Receiver Operating Characteristic (auROC) are selected. These two metrics are best indicators of the model performance in an unbalanced scenario.

## III. results

### A. Selecting the best model

In order to evaluate the three parameters of our interest: length of the input sequence, type of embedding, and different predictive model, we implemented 30 different predictive models in which one is changing. All of these models were implemented in python with scikit-learn and keras libraries for SVM and CNN, respectively. All hyperparameters for SVM and CNN models were tuned with grid search. The details of these parameters are stated in the codes provided in the github repository of this paper. The results based on auPR are summarized in Table 1.

**Table 1:** Comparison of different input sequence lengths, embedding methods and predictive models on auPR of testing samples.

|  |  | SVM |  |  | CNN |  |  |
| --- | --- | --- | --- | --- | --- | --- | --- |
| Input length (L) |  | One-hot | dna2vec |  | One-hot | dna2vec |  |
|  |  |  | K = 3 | K = 5 |  | K = 3 | K = 5 |
| **2'O**<br>**in center** | 45 nc | 0.53 | 0.66 | 0.63 | **0.78** | 0.74 | 0.70 |
|  | 75 nc | 0.55 | 0.66 | 0.63 | 0.74 | 0.75 | 0.70 |
|  | 105 nc | 0.52 | 0.61 | 0.63 | 0.76 | 0.75 | 0.71 |
| **2'O +**<br>**44 nc** | Up | 0.47 | 0.50 | 0.48 | 0.46 | 0.50 | 0.52 |
|  | Down | 0.45 | 0.53 | 0.55 | 0.49 | 0.52 | 0.63 |

As can be observed in Table 1, the best model performance was achieved when one-hot encoding was used with inputs with a length of 45 nc in the CNN model. Among SVM models, model with input length 45 nc and dna2vec (kmer=3) achieved the best performance. In **Figure 2** (A) and **Figure 2**(B) PR and ROC curves of 5 fold cross-validation of these two models are shown.

**Figure 2:**
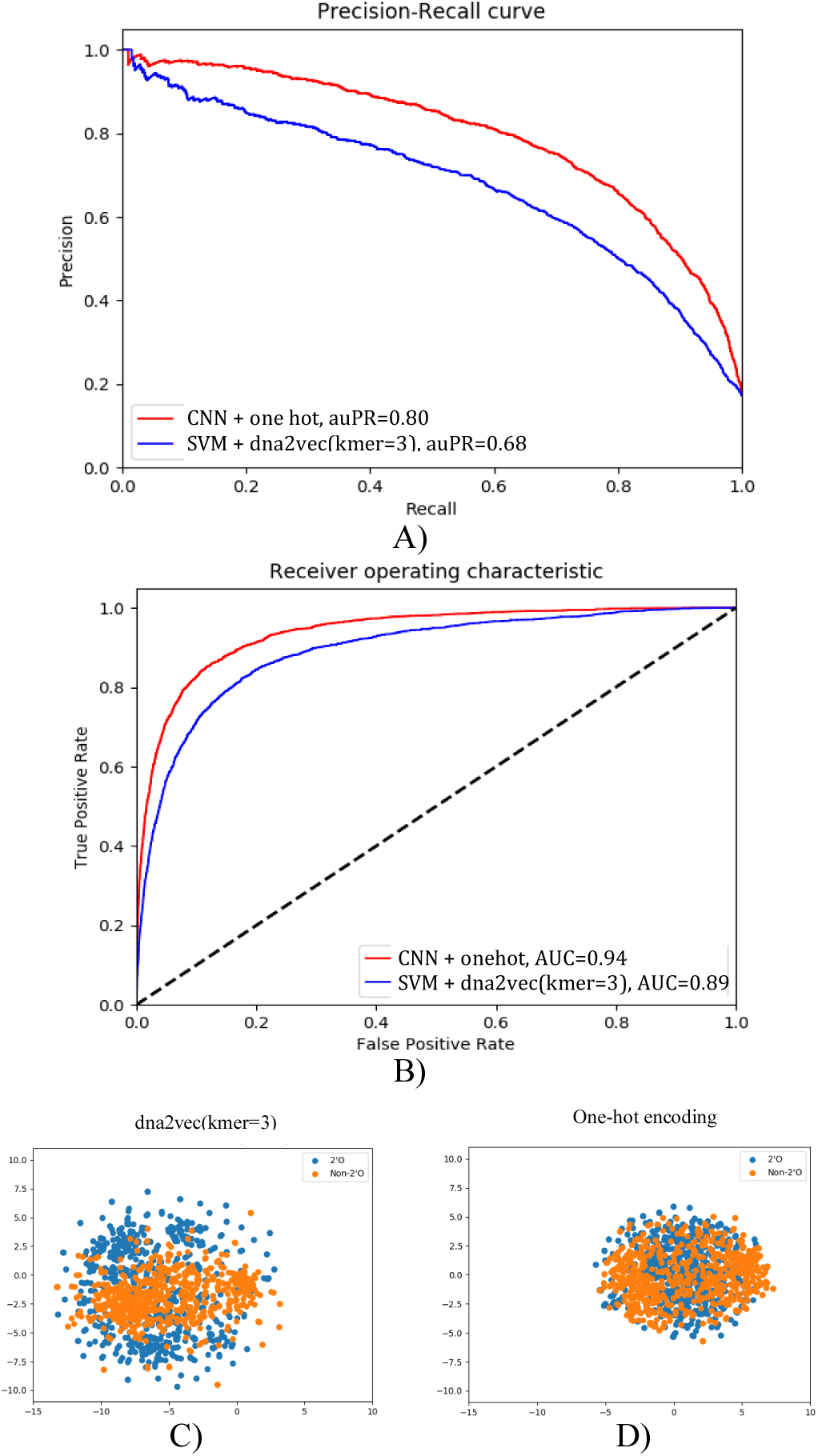
5 fold cross-validation of best SVM and CNN models. A) Precision-Recall curve B) Receiver operating characteristic. C) t-SNE plot of input sequences with length 45 nc for dna2vec D) one-hot encoding.

There are several interesting observations that clarify the impact of each parameter in our predictions. Firstly, we can claim that increasing the length of input sequences does not improve the performance of models. In addition, when models are trained with 2′O sites with their 44 downstream nc, their performance is slightly better. This means there are existing patterns on both sides of 2′O sites that enable models to have fair prediction results, but the downstream leads to higher results, which might be caused by N_m_ motif.

Moreover, different embedding methods create a distinguishable difference between the performance of SVM models. This provides evidence that ML models are highly dependent on the feature selection method and why many research studies have attempted to find the best feature selection method. On the contrary, when CNN is utilized, the performance of the one-hot encoding is better than dna2vec. This elucidates that although dna2vec is providing richer features in SVM models, it does not help to improve the CNN model, which has its own automatic feature selection method. In addition, changing kmer from three to five does not have any effects on predictive models. In order to have a better understanding of these embeddings, t-SNE plot of 600 held out test samples with one-hot and dna2vec (with kmer=3) for input length of 45 nc are plotted in **Figure 2**(A) and **Figure 2**(B). As can be seen in the t-SNE plots, one-hot encoding has a more condensed representation than dna2vec, which might be the reason it yields lower performance in SVMs. To make further analysis on model interpretation, we select the CNN model with an input length of 45 nc (2′O centered in the middle) and the one-hot encoding for the investigation of the next sections.

### B. Significant nucleotides around 2′O sites

In this section, we utilize the Integrated Gradient (IG) method in [14], a visualization method that aims to interpret DL models with respect to their inputs, to understand the impact of each nucleotide from a model perspective. Our main goal is to find out which of the nucleotides and at what distance from 2′O site has the greatest and lowest influence in the predictions. All of the 2802 2′O samples with surrounding nucleotides (i.e. 22 from downstream and upstream) were used to generate significance scores at each nucleotide of input. We averaged the values yielded from IG method for all the positive samples in each position and finally resulted in one matrix with length 4*45, same as the input dimension. In **Figure 3**, the attribution scores corresponding to every nucleotide is plotted. As it is shown in **Figure 3**, an RNA sequence with length 45 nucleotides where 2′O site is centered in the middle (i.e. position 23). The scores vary from [-1, 1] for every nucleotide at each location where positive and negative values are indicators of a positive and negative effect in predictions, respectively.

**Figure 3:**
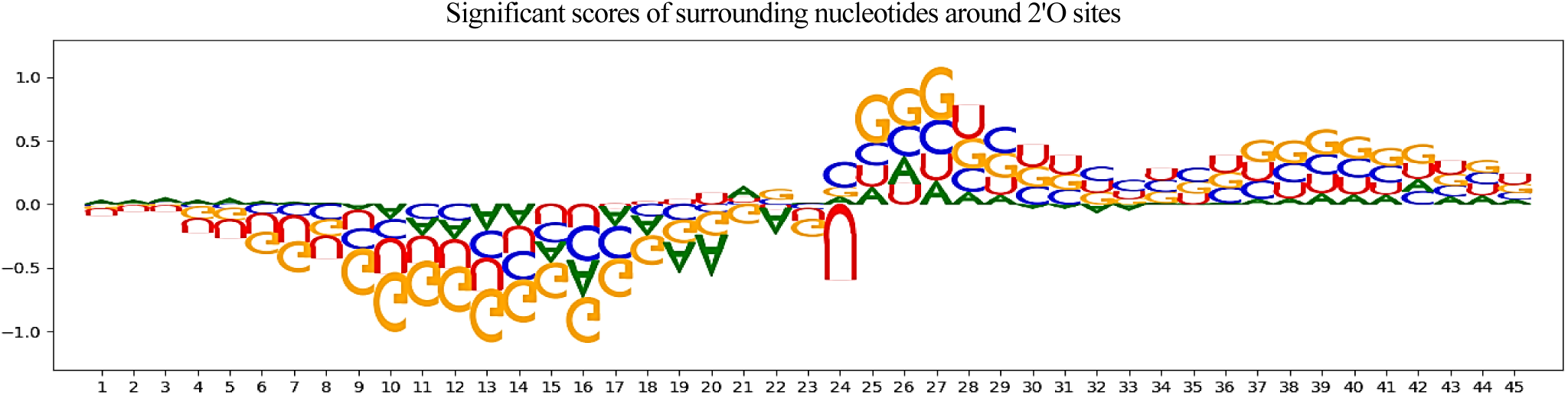
Significant nucleotides around 2′O (in 23^rd^ position) through IG method with CNN + one hot + input length 45 nc.

Several interesting phenomena can be observed in this significant score matrix. Firstly, the model pays the highest attention at the location of motif N_m_AGAUCGAGGA that was founded in [7]. The length of the significant score on the downstream of 2′O is as big as the length of the motif that was reported. This fact illustrates that the CNN model is mainly looking at existing motif in the 2′O sequences. In addition, significant negative scores on the upstream of 2′O sites do not mean these nucleotides have no impact on the prediction results. Indeed, as explained in [15], they have a lower contribution effect in predictions.

## IV. Discussion and conclusion

In this paper, we presented a comprehensive investigation of building ML and DL models for predicting 2′O locations in RNA sequences. Specifically, three critical factors in building the predictive models namely, the length of the input sequence containing 2′O site, the embedding method, and the type of predictive model were studied individually. SVM and CNN were selected as the best predictive models among ML and DL models, respectively.

According to our predictive models, embedding can be an influential parameter in the performance of SVM models. However, it does not ameliorate the performance of CNN models. Moreover, it was shown that models with downstream and upstream of 2′O sites can be used for building a predictive model. Due to the existence of a dominant 2′O motif in downstream of inputs, models would have better performance when downstream is utilized. Furthermore, interpretation methods were harnessed to further explore the importance of surrounding nucleotides around 2′O sites. It was shown that the model pays the highest attention to the downstream of RNA sequence for predictions that further elucidate the impact of the dominant motif in the downstream of 2′O sites.

deeplift [15], an improved version of IG, was also used to investigate the significant score matrix. However, we found this method to be highly dependent on the required reference sequence. Besides, the amplitude of significant scores in each location can be further argued. Since IG is based on a numerical stochastic method that requires initialization, the amplitudes can vary slightly if the model is re-trained. Although amplitudes of values slightly change, the location of significant scores remains the same after re-training of the model. This is a new and active research area that still requires more studies to solve such problems.

